# A comprehensive germline variant and expression analyses of *ACE2*, *TMPRSS2* and SARS-CoV-2 activator *FURIN* genes from the Middle East: Combating SARS-CoV-2 with precision medicine

**DOI:** 10.1101/2020.05.16.099176

**Authors:** Fahd Al-Mulla, Anwar Mohammad, Ashraf Al Madhoun, Dania Haddad, Hamad Ali, Muthukrishnan Eaaswarkhanth, Sumi Elsa John, Rasheeba Nizam, Arshad Channanath, Mohamed Abu-Farha, Rasheed Ahmad, Jehad Abubaker, Thangavel Alphonse Thanaraj

**Author notes:** **Corresponding author:** Prof. Fahd Al-Mulla, Dasman Diabetes Institute, P.O. Box 1180, Dasman 15462, Kuwait.

## Abstract

The severity of the new COVID-19 pandemic caused by the SARS-CoV-2 virus is strikingly variable in different global populations. SARS-CoV-2 uses *ACE2* as a cell receptor, *TMPRSS2* protease, and *FURIN* peptidase to invade human cells. Here, we investigated 1,378 whole-exome sequences of individuals from the Middle Eastern populations (Kuwait, Qatar, and Iran) to explore natural variations in the *ACE2*, *TMPRSS2,* and *FURIN* genes. We identified two activating variants (K26R and N720D) in the *ACE2* gene that are more common in Europeans than in the Middle Eastern, East Asian, and African populations. We postulate that K26R can activate *ACE2* and facilitate binding to S-protein RBD while N720D enhances *TMPRSS2* cutting and, ultimately, viral entry. We also detected deleterious variants in *FURIN* that are frequent in the Middle Eastern but not in the European populations. This study highlights specific genetic variations in the *ACE2* and *FURIN* genes that may explain SARS-CoV-2 clinical disparity. We showed structural evidence of the functionality of these activating variants that increase the SARS-CoV-2 aggressiveness. Finally, our data illustrate a significant correlation between *ACE2* variants identified in people from Middle Eastern origins that can be further explored to explain the variation in COVID-19 infection and mortality rates globally.

## Introduction

The global pandemic COVID-19 caused by SARS-CoV-2 virus is life-threatening and has become a significant concern to humanity. Notably, the severity of this disease is highly variable in different populations across the world ^1^. Since the outbreak, several studies have reported specific factors including age, gender, and pre-existing health conditions that could have contributed to the increased severity of the disease ^2–4^. The genetic susceptibility to COVID-19 has also been explored by scrutinizing Angiotensin converting enzyme 2 (*ACE2*) genetic variations in different populations. *ACE2* is the functional receptor mediating entry of SARS-CoV-2 into the host cells ^5^, which is facilitated by *FURIN* cleavage ^6–8^. Transmembrane serine protease 2 (*TMPRSS2*) is another candidate gene that has been linked to COVID-19 disease ^8–10^. *TMPRSS2* expression enhances *ACE2*-mediated SARS-CoV-2 cell invasion by operating as a co-receptor ^10^. The increased cleavage activity of this protease was suggested to diminish viral recognition by neutralizing antibodies and by activating SARS spike (S) protein for virus-cell fusion ^11^ and facilitates the active binding of SARS-CoV-2 through *ACE2* receptor, which is a risk factor for a more serious COVID-19 presentation ^8–10^.

The hallmark of the novel SARS-CoV-2, as compared to other SARS viruses, is the presence of a polybasic *FURIN* cleavage site. *FURIN* has been reported to facilitate the transport of SARS-CoV-2 into or from the host cell ^6–8^. Notably, a recent study has highlighted the presence of a unique functional polybasic *FURIN* cleavage consensus site between the two spike subunits S1 and S2 by the insertion of 12 nucleotides encoding PRRA in the S protein of SARS-CoV-2 virus ^12^. The *FURIN*-like cleavage-site is cleaved during virus egress, which primes the S-protein providing a gain-of-function for the efficient spreading of the SARS-CoV-2 among humans ^13,14^. It is, therefore, likely that the presence of a deleterious *ACE2*, *TMPRSS2* and *FURIN* gene variants may modulate viral infectivity among humans, making some people less vulnerable than others. Recent studies, assessed the genetic variations and eQTL (expression quantitative trait locus) expression profiles in the candidate genes *ACE2*, *TMPRSS2*, and *FURIN* to demonstrate the sex and population-wise differences that may influence the pathogenicity of SARS-CoV-2 ^9,15–19^. It is to be noted that these studies focused only on the European and East Asian populations. Given the extremely high prevalence of obesity (80%), hypertension (28%) and diabetes (20%) of the population in the Gulf states ^20–22^ which are considered as risk factors for mortality from COVID-19 ^4,23^, the witnessed low infectivity and mortality rates registered in this area of the world are intriguing. Even though this could be due to various factors that are not well accounted for yet such as testing, hot weather or extreme measures taken early on by some countries, it could also be due to ethnic genetic variations in the *ACE2*-*TMPRSS2*-*FURIN* genes that are key regulators for orchestrating SARS-CoV-2 cellular access.

As a result, it is crucial to study the variation of these candidate genes *ACE2*, *TMPRSS2* and *FURIN* in Middle Eastern populations to better understand possible natural genetic components that can be responsible for these differences. Here, we present a comprehensive comparative assessment of deleterious or gain of function mutations of *ACE2*, *TMPRSS2,* and *FURIN* that may have influenced disease progression in the Middle Eastern populations compared to European, African and East Asian populations. While these findings are preliminary and based on our genetic data and worldwide public datasets, they present a compelling perception into the role of naturally occurring genetic variants in *ACE2*, *TMPRSS2,* and *FURIN* in different populations. They can shed light on the reported variations in susceptibility or resistance to SARS-CoV-2 infection in different populations and can be available for other scientists to utilize them in precision medicine.

## Results

### *ACE2* receptor variations in the Middle East

We first examined *ACE2* gene variation frequencies in Kuwait, Qatar, East Asia, and Africa where the impact of SARS-CoV-2 has been modest and compared it to Iran, which is moderately affected and then to Europe, the continent with the most deaths per population. Overall, we found human *ACE2* gene variations and the probability of loss of function mutations (pLOF=0.1, C.I. 0.04-0.25) to be low in comparison to *ACE* (pLOF=0.87, C.I. 0.71-1.08), which is a gene of similar size, indicating that the *ACE2* gene is highly intolerant to loss of function mutations. Additionally, we identified 19 missense variants in the *ACE2* gene from Kuwait, Qatar and Iran (Table 1). All were rare variants defined by minor allele frequency (MAF) of less than 1%. The genetic variants included four novel variants from Kuwait, and six from Iran.

**Table 1.**
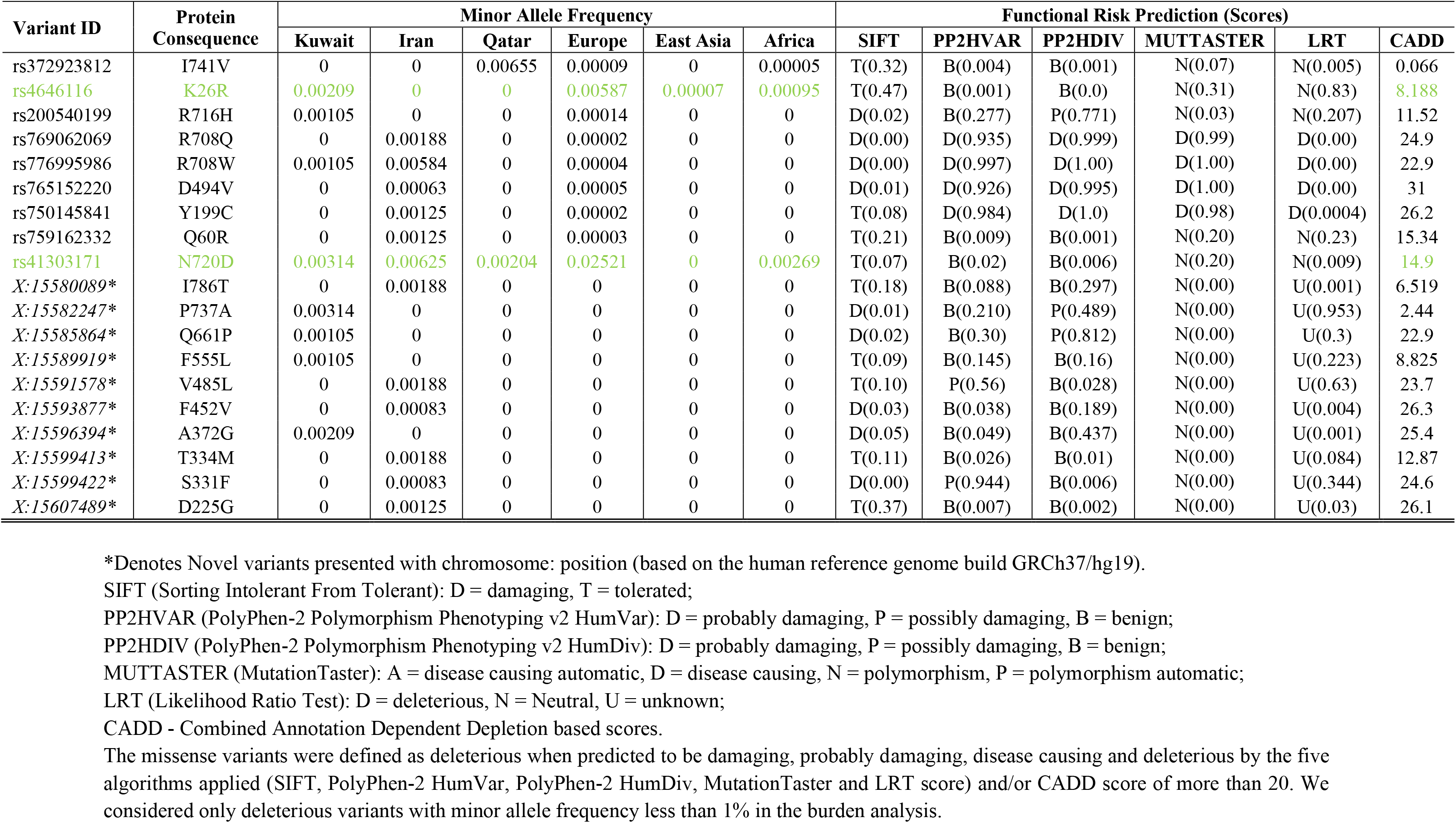
*ACE2* missense variants in the Middle East and gnomAD populations.

We identified four deleterious variants (rs776995986, rs769062069, rs765152220, and rs750145841), causing R708W, R708Q, D494V, Y199C missense amino acid substitutions in the *ACE2* gene using risk prediction tools as described in the methods section (Table 1). All the *ACE2* gene deleterious variants were absent from the African, East Asian and Qatari data and were very rare in Europeans but were present at MAF of 0.063-0.5% in the Iranian population (Table 1). This suggests a more protective effect and a significant decrease in the disease burden in Iran compared to Europe (p<0.05; Table 2). The positions of the *ACE2* receptor polymorphisms on the linearized *ACE2* protein model are shown in Supplementary Fig. 1, and 3D-models for the same are shown in Supplementary Fig. 2 and Supplementary Fig. 3. It is noteworthy that none of the *ACE2* polymorphisms identified in this study involved the three *ACE2* regions known to directly bind the SARS-CoV-2 S-Protein Receptor Binding Domain (RBD), namely amino acids 30-41, 82-84, and 353-357 (Supplementary Fig. 1).

**Table 2.** Burden of *ACE2* rare variants in the Middle East and gnomAD populations.

| Variant ID | Protein Consequence | Minor Allele Frequency |  |  |  |  |  | P value |  |  |
| --- | --- | --- | --- | --- | --- | --- | --- | --- | --- | --- |
|  |  | KWT | IRN | QAR | EUR | EAS | AFR | KWT vs IRN | KWT vs EUR | IRN vs EUR |
| rs776995986 | R708W | 1.05E-03 | 5.64E-03 | 0.00 | 3.79E-05 | 0.00 | 0.00 | 0.1084 | 0.0515 | 0.0005 |
| rs769062069 | R708Q | 0.00 | 1.88E-03 | 0.00 | 1.46E-05 | 0.00 | 0.00 | — | — | 0.0005 |
| rs765152220 | D494V | 0.00 | 6.25E-04 | 0.00 | 5.37E-05 | 0.00 | 0.00 | — | — | 0.0995 |
| rs750145841 | Y199C | 0.00 | 1.25E-03 | 0.00 | 2.00E-05 | 0.00 | 0.00 | — | — | 0.0015 |

The missense variants were defined as deleterious when predicted to be damaging, probably damaging, disease causing and deleterious by the five algorithms applied (SIFT, PolyPhen-2 HumVar, PolyPhen-2 HumDiv, MutationTaster and LRT score) and/or CADD score of more than 20. We considered only deleterious variants with minor allele frequency less than 1% in the burden analysis. P values are calculated using Chi-square test. KWT-Kuwaitis; IRN-Iranians; QAR-Qataris; EUR-Europeans (non-Finnish); EAS-East Asians; AFR-Africans.

Next, we examined whether natural *ACE2* gene variations that increase the affinity of *ACE2* to the S-protein or facilitate viral entry/viral load exist more frequently in high-burden compared to low-burden populations. Two such genetic variants existed in our data (Table 1). The first, rs4646116, is a missense variant that changes a lysine amino acid at position 26 to arginine (K26R) (Table 1). The K26, which is just proximal to the first region of the *ACE2* receptor involved in S-protein binding, has been shown previously to bind the sterically hindering first mannose in the glycan that is linked to N90 and thus stabilizes the glycan moiety hindering the binding of S-protein RBD to *ACE2* ^24^ (Fig. 1a). The missense variant R26 creates a new hydrogen bond with D30, which is then poised to build a salt-bridge with the S-protein RBD K417 that increases the affinity of SARS-CoV-2 to the *ACE2* receptor ^18^ (Fig. 1b). Indeed, the *ACE2* K26R activating variant was extremely rare in East Asian (MAF=0.007%), Africans (MAF=0.095%), but the second most common variant in Europeans with MAF of 0.587% (shown in green fonts in Table 1). The MAF of this variant in the Kuwaiti population was nearly half that of Europeans (MAF=0.29%), and it was absent from the Qatari and Iranian exome data (Table 1). Our structural modeling supports the notion that K26R is an *ACE2* receptor activating variant (Fig. 1a, b). Consistent with these findings, using a synthetic human *ACE2* mutant library, a recent study reported that the R26 variant increased S-protein binding and susceptibility to the virus significantly ^25^.

**Figure 1.**
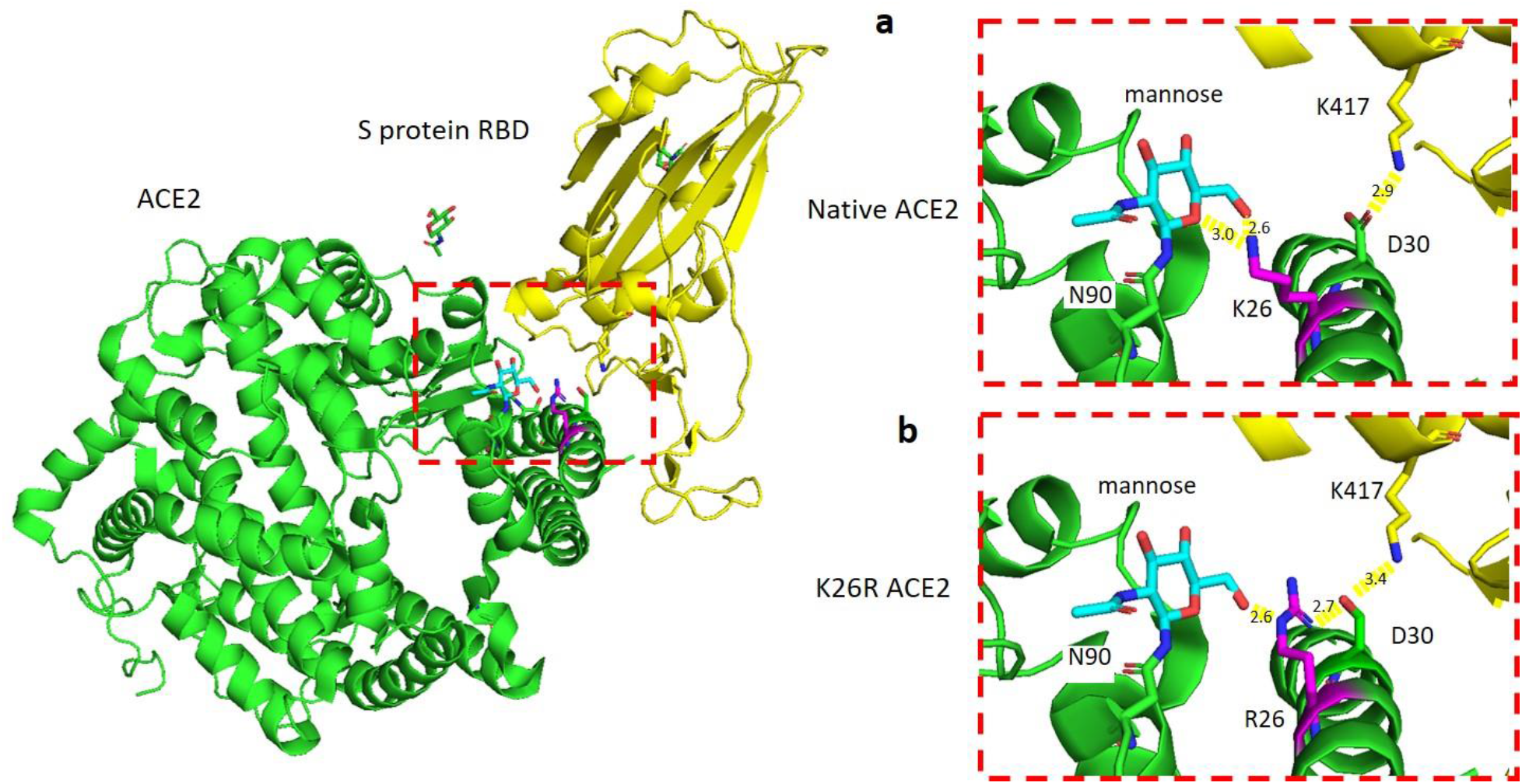
K26R polymorphism in *ACE2* mapped to the structure of *ACE2*, green; in complex with S protein RBD, yellow (PDB ID: 6VW1). **(a)** In native *ACE2*, K26, magenta, forms two H-bonds with the mannose moiety, cyan, of *ACE2* N90 which may stabilize the glycan-*ACE2* complex. **(b)** The mutant R26, magenta, forms one H-bond with mannose. A process that could provoke the moiety-*ACE2* stability and increases the affinity of the *ACE2* a-helix to S protein RBD binding, where, R26 functions as a backbone and interacts with D30 which in turn dignified to build a salt-bridge with the S protein RBD K417, yellow stick.

We also subjected novel and known *ACE2* variants to structural predictions that may impact the binding of SARS-CoV-2 to the host cells. These changes (A372G, Y199C, D225G, F452V and V485L) were located proximal to the protein residues that mediate its activity. A previous study indicated that the three amino acid (aa) regions 30-41, especially the residue near lysine 31 and tyrosine 41, 82-84 and 353-357 in *ACE2* were essential for the binding of S-protein in coronavirus ^26^. Our structural estimation based on the 19 *ACE2* missense variants identified in Kuwaitis, Qataris and Iranians (Table 1), revealed the location of K26R variant near the aa residues 30-41 and the novel variants, S331F and T334M, adjacent to tyrosine at 353-357. The A372G is adjacent to the zinc-binding domain (HEMGH 374-378) critical for *ACE2* enzymatic activity ^27^. While Y199C, D225G, F452V, and V485L all fall within the metallocarboxypeptidase domain (aa 18-617) ^28^ (Supplementary Fig. 1).

### *ACE2* N720D receptor variation and structural modeling

The second activating, and by far the most common *ACE2* gene variant in Europeans (MAF=2.52%) and Italians (MAF=1.6%; detected in 105 of 6984 exomes) ^17^ was rs41303171, which replaces the amino acid asparagine at position 720 to aspartic acid (N720D) (Green font in Table 1). This *ACE2* variant was absent from the East Asian population (13,784 exomes) and was significantly rarer in the Middle East and Africa (Table 1). This particular variant has been reported before, but its clinical relevance was persistently dismissed because the codon 720, being far from *ACE2*-Spike protein interface, does not appear to be an obvious candidate for *ACE2* receptor binding to the S-protein of SARS-CoV-2 ^18,29^. However, we noted that N720D is located 4 amino acids proximal to the *TMPRSS2* cleavage site (aa 697-716) as shown in Supplementary Fig. 1. Recent studies demonstrated that *TMPRSS2* cleavage of the *ACE2* receptor increases SARS-CoV-2 cellular entry ^30,31^. It is not unreasonable to suggest that this activating variant may play a similar role in SARS-CoV-2, rendering people who harbor it more prone to severe infection and higher viral load.

*ACE2* cleavage by *TMPRSS2* enhances the S-protein viral entry ^5^. *TMPRSS2* cleaves *ACE2* between residues 697 and 716, which is the third and fourth helices in the C-terminal collectrin neck domain of the dimer interface of *ACE2* ^30^. To further dissect the mechanisms underlying the enhanced *ACE2*-*TMPRSS2* accessibility, we performed structural analysis of the recently published *ACE2* domain bound to B0AT1 (PDB ID:6M18) ^5^ and showed that N720 is located on the same interface of the loop region in close vicinity to the *TMPRSS2* cleavage site (Fig. 2a). Since loop region of protein are unordered and display conformational dynamics, a mutation close to the cleavage site can affect the binding affinity of *TMPRSS2*. Therefore, we used DynaMut web server to predict the effect of mutation N720D on the stability and flexibility of *ACE2* ^32,33^. Whereby, the predicted stability change was (ΔΔG): −0.470 Kcal/mol, which indicated the destabilization of *ACE2* receptor after the introduction of D720 (Fig. 2b). The N720D mutation has resulted in an increase in entropy in the loop region (ΔΔSVib: 0.070 kcal.mol^−1^.K^−1^) ^33^ (Fig. 2b), depicting a more unstable state, which makes *ACE2* more readily cleavable by *TMPRSS2*.

**Figure 2.**
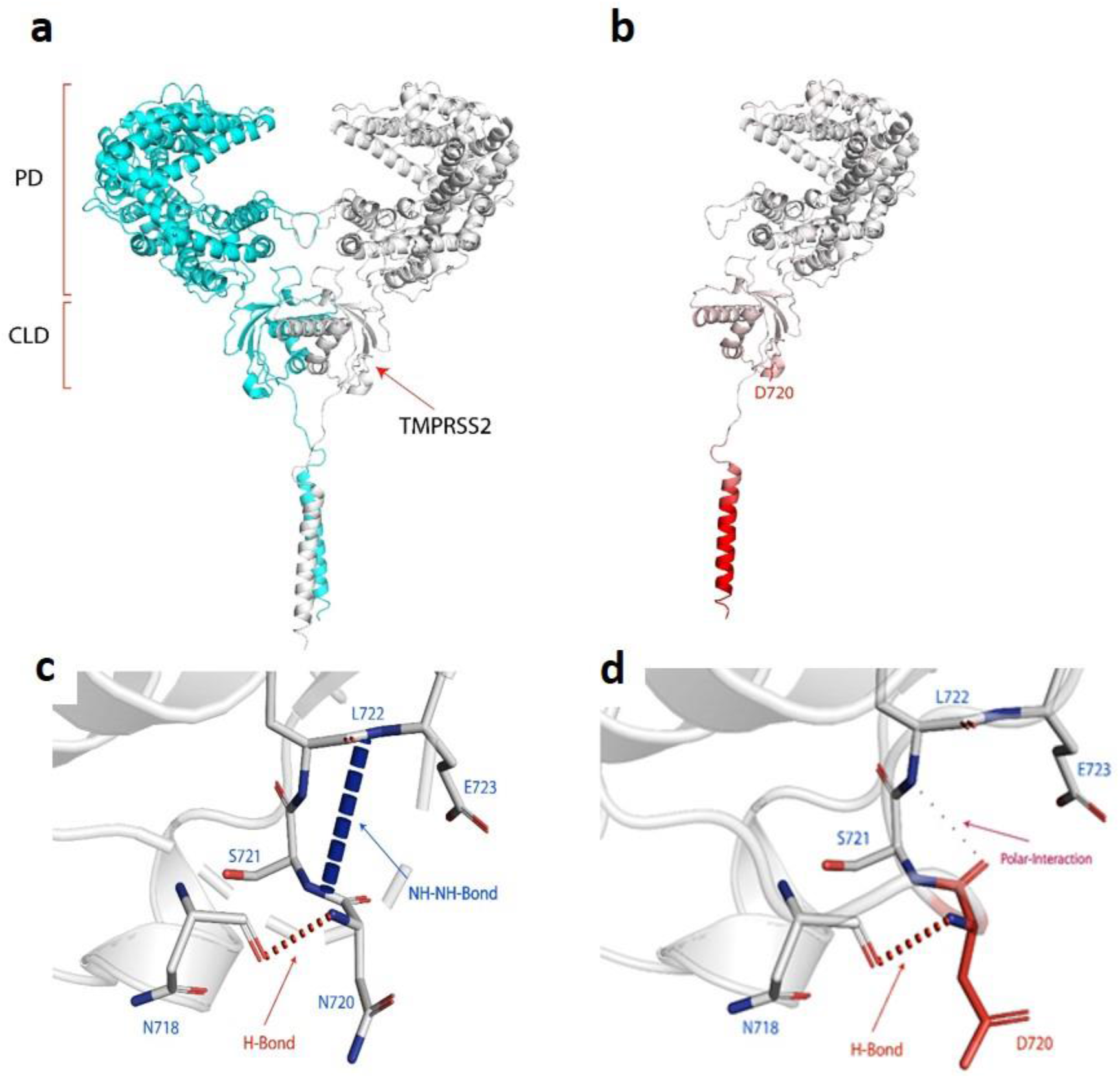
**(a)** *ACE2* dimer complex (promoter 1 Blue), (promoter 2 white). The peptidase domain (PD) is the site where the SARS-CoV-2 S protein ribosomal domain (RBD) binds. The C-terminal collectrin-like domain (CLD) is the region where the *TMPRSS2* cleaves *ACE2*. **(b)** Single domain of the *ACE2* mutation N720D, the red region of the protein depicted the more flexible region of the protein due to the N720D mutation with a (ΔΔG): −0.470 Kcal/mol and ΔΔS: 0.070 kcal.mol^−1^.K^−1^ (increase of molecule flexibility). **(c)** Depicts N720 backbone NH forming a H-bond with N718 and E723 forming an NH-NH bond with S721. **(d)** D720 backbone NH forming a H-bond with N718 and COO-group of D720 forming a weak polar interaction with L722 backbone.

In addition, we modelled in the various non-covalent interactions for both N720 and mutant D720 and other amino acids in the loop region (Fig. 2c, d). In Fig. 2c, N720 formed a backbone hydrogen-bond with N718, this conformation also resulted in an NH-NH between E723 and S721. Whereas, with the activating variant D720 (Fig. 2d) the back-bone hydrogen bond with N718 is still established, however, the conformational change has resulted in a polar interaction between the backbone D720 COO-group and backbone NH of L722, forming a weak polar interaction (through water-mediated hydrogen bond). As such, the D720 variant altered the conformation of the loop, and the NH-NH bond between E723 and S721 cannot be formed. Whereby, such intermolecular amide interactions are significant for protein stability ^34^. Such a change in interatomic interactions between amino acids near the cleavage region can decrease the stability and increase the flexibility of the loop ^35^, which makes it easier for *TMPRSS2* to cleave.

To gain insights that are pathologically and clinically relevant, we then asked whether a correlation exists between the N720D MAF data and mortality rates reported in corresponding regions. We identified a significant correlation between N720D MAF and deaths per 1 million reported from different regions (Fig. 3) with Pearson’s correlation coefficient of 99.4% (p<0.0001; C.I. 0.945-0.99).

**Figure 3.**
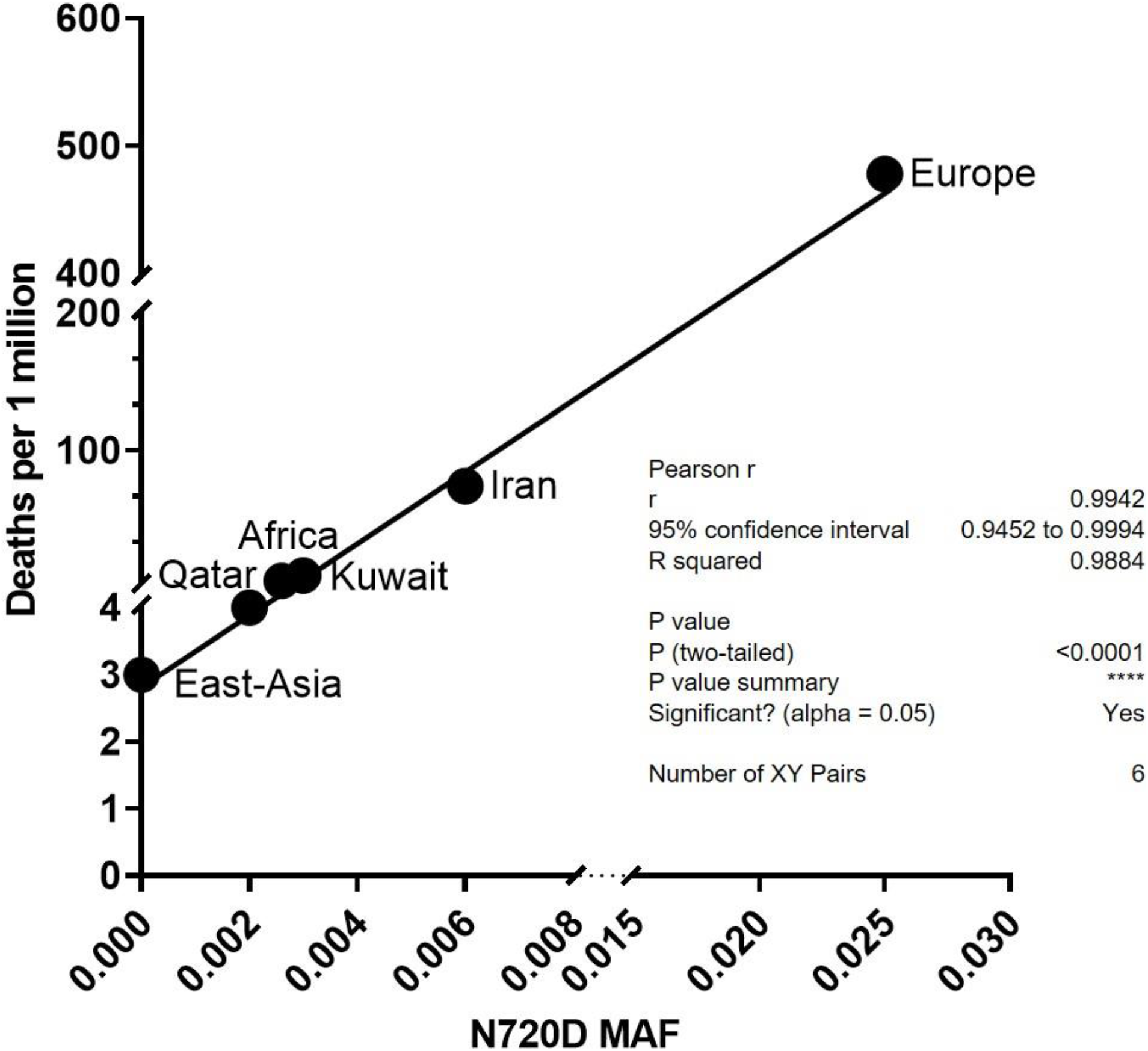
The plot shows the significant correlation between N720D minor allele frequency (MAF) and deaths per 1 million reported from different regions with Pearson’s correlation coefficient of 99.4% (p<0.0001; C.I. 0.945-0.99).

Next, we examined *ACE2* eQTL variants in the Middle Eastern populations (Supplementary Table 1). There were few variants in the Kuwaiti and Iranian populations that influenced *ACE2* expression. In the Qatari population the variants modifying *ACE2* gene expression were similar in frequencies to Europeans (Supplementary Table 1) with most upregulating *ACE2* expression in the brain and tibial nerve tissues (https://gtexportal.org/home/). Notably, the tibial nerves were affected in diabetic conditions ^36^, which inflicts a high number of patients in the Middle East. The most downregulating eQTL variant, rs112171234, was present in 20% and 6% of the African and Qatari populations respectively. Overall, the eQTL data pertaining to *ACE2* expression were not significantly predictive nor informative.

### *TMPRSS2* Variation

Five rare and one common deleterious variants were identified in the *TMPRSS2* gene in the Middle Eastern population (Supplementary Fig. 4, Supplementary Table 2, Supplementary Table 3). Overall, there was no significant conclusion that may be withdrawn from the *TMPRSS2* genetic variation data. On the expression level, we discerned four *TMPRSS2* eQTL variants, which were detected only in Qataris among the Middle Eastern populations (Supplementary Table 4). One of these variants, rs6517673 is intronic and downregulates the expression of *TMPRSS2* in the prostate. Two of the eQTL variants, rs79391937 and rs79566442, decrease the expression of the *TMPRSS2* gene in the thyroid and ovary tissues, respectively. While the variant rs11701542 upregulates the *TMPRSS2* gene expression in the testis (https://gtexportal.org/home/). It is worth noting that two-third of the mortality due to COVID-19 disease affects males ^15,37^.

### *FURIN* Variation

Genetic variation analysis of the *FURIN* gene resulted in the identification of 13 known missense variants (Table 3, Supplementary Fig. 5). Like *ACE2*, all the identified variants were rare in the Middle Eastern populations. However, unlike the *ACE2* gene, no novel variants were observed in *FURIN* in the Middle Eastern population (Table 3). Among the 13, we detected seven deleterious variants suggesting a possible decrease in *FURIN* protease function, which can potentially reduce the risk of SARS-CoV-2 in the studied populations. In this context, deleterious *FURIN* gene variations were observed least in East Asians, Africans, followed by Europeans then Iranians (Table 4). Interestingly, both in Qatar and Kuwait the deleterious *FURIN* genetic variations were more common, suggesting a possible protective effect against the SARS-CoV-2 (p<0.05; Table 4). For example, the MAF of R37C, and R81C in Kuwait and Qatar were 0.4%/0.7% and 2.4%/3.4% respectively compared to significantly lower MAF in corresponding variants in Europeans (p<0.05; Table 4). Together, these data support the premise that *FURIN* gene variants may play important roles in protection against SARS-CoV-2 in the Middle East. It is worth noting, however, that in Africa *FURIN* gene variants may not be a contributing factor in viral protection (Table 4). We next sought to determine the extent of *FURIN* expression in the Middle East. We detected 16 *FURIN* eQTL variants in the Middle Eastern populations, most of which were reported in the Qatar population (Supplementary Table 5). We observed a high frequency of the *FURIN* upregulating variants, rs6226 (93%) and rs8039305 (81%) in the African populations compared to the Middle Eastern, European and East Asian populations (p<0.05) (Supplementary Table 5). In the GeneATLAS PheWAS database ^38^, these frequent variants were significantly related to hypertension (rs6226: P=1.3569e-09, OR=1.03; rs8039305: P= 1.7643e-38, OR=1.05), which was one of the common comorbid conditions associated with the increased risk of SARS-CoV-2 infection in Chinese ^2,3^ and American ^39^ populations.

**Table 3.** *FURIN* known missense variants in the Middle East and gnomAD populations.

| Variant ID | Protein Consequence | Minor Allele Frequency |  |  |  |  |  | Functional Risk Prediction (Scores) |  |  |  |  |  |
| --- | --- | --- | --- | --- | --- | --- | --- | --- | --- | --- | --- | --- | --- |
|  |  | Kuwait | Iran | Qatar | Europe | East Asia | Africa | SIFT | PP2HVAR | PP2HDIV | MUTTASTER | LRT | CADD |
| rs201172453 | R37C | 0.00418 | 0.00375 | 0.00700 | 0.00007 | 0 | 0.00016 | D (0.01) | P (0.586) | D (0.995) | N (0.000) | N (0.052) | 22.7 |
| rs749761700 | R37H | 0.00105 | 0 | 0 | 0.00001 | 0 | 0 | T (1.00) | B (0.001) | B (0.001) | N (0.000) | N (0.052) | 0.496 |
| rs16944971 | A43V | 0.00628 | 0.00188 | 0.00500 | 0.00255 | 0 | 0.07835 | T (0.25) | B (0.007) | B (0.010) | P (0.411) | N (0.007) | 11.82 |
| rs148110342 | R81C | 0.02406 | 0.00750 | 0.03400 | 0.00159 | 0 | 0.00016 | T (0.17) | P (0.586) | D (0.989) | D (0.995) | N (0.051) | 23 |
| rs751909359 | R86Q | 0 | 0.00070 | 0 | 0.00003 | 0 | 0 | T (0.31) | B (0.028) | B (0.244) | D (0.999) | D (0.000) | 22.3 |
| rs116359616 | V109M | 0.00105 | 0 | 0 | 0.00002 | 0 | 0.02318 | D (0.03) | B (0.010) | B (0.083) | D (0.635) | N (0.168) | 4.625 |
| rs143276283 | Q399R | 0.00209 | 0.00063 | 0 | 0.00006 | 0 | 0 | T (1.00) | B (0.002) | B (0.000) | D (0.967) | N (0.002) | 18.69 |
| rs750446344 | V580I | 0.00314 | 0 | 0 | 0.00004 | 0 | 0 | T (0.23) | B (0.007) | B (0.010) | N (0.000) | N (0.835) | 0.211 |
| rs752639409 | R637Q | 0 | 0.00070 | 0 | 0 | 0 | 0.00006 | T (0.58) | B (0.051) | B (0.200) | N (0.003) | N (0.031) | 21.3 |
| rs761541008 | R677W | 0 | 0.00070 | 0 | 0.00005 | 0 | 0 | T (0.05) | B (0.332) | P (0.947) | N (0.046) | D (0.000) | 27.4 |
| rs193268286 | S685P | 0 | 0 | 0 | 0.00004 | 0 | 0 | T (0.09) | D (0.994) | D (0.999) | D (0.999) | D (0.000) | 26.6 |
| rs760368022 | H712Y | 0.00105 | 0 | 0 | 0.00001 | 0 | 0 | D (0.01) | B (0.089) | B (0.367) | N (0.016) | N (0.032) | 19.66 |
| rs35641241 | R745Q | 0 | 0 | 0 | 0.00118 | 0.00102 | 0.00008 | T (0.13) | B (0.375) | D (0.989) | N (0.149) | N (0.021) | 23.4 |

SIFT (Sorting Intolerant From Tolerant): D = damaging, T = tolerated;
PP2HVAR (PolyPhen-2 Polymorphism Phenotyping v2 HumVar): D = probably damaging, P = possibly damaging, B = benign;
PP2HDIV (PolyPhen-2 Polymorphism Phenotyping v2 HumDiv): D = probably damaging, P = possibly damaging, B = benign;
MUTTASTER (MutationTaster): A = disease causing automatic, D = disease causing, N = polymorphism, P = polymorphism automatic;
LRT (Likelihood Ratio Test): D = deleterious, N = Neutral, U = unknown;
CADD - Combined Annotation Dependent Depletion based scores.
The missense variants were defined as deleterious when predicted to be damaging, probably damaging, disease causing and deleterious by the five algorithms applied (SIFT, PolyPhen-2 HumVar, PolyPhen-2 HumDiv, MutationTaster and LRT score) and/or CADD score of more than 20. We considered only deleterious variants with minor allele frequency less than 1% in the burden analysis.

**Table 4.** Burden of *FURIN* rare variants in the Middle East and gnomAD populations.

| Variant ID | Protein Consequence | Minor Allele Frequency |  |  |  |  |  | P value |  |  |  |  |  |  |  |  |  |  |  |
| --- | --- | --- | --- | --- | --- | --- | --- | --- | --- | --- | --- | --- | --- | --- | --- | --- | --- | --- | --- |
|  |  | KWT | IRN | QAR | EUR | EAS | AFR | KWT vs IRN | KWT vs QAR | KWT vs EUR | KWT vs AFR | IRN vs QAR | IRN vs EUR | IRN vs AFR | QAR vs EUR | QAR vs AFR | EUR vs EAS | EUR vs AFR | EAS vs AFR |
| rs201172453 | R37C | 0.004 | 0.004 | 0.007 | 7.0E-05 | 0 | 1.6E-04 | 1 | 1 | 0.0005 | 0.0005 | 1 | 0.0005 | 0.0005 | 0.0120 | 0.0610 | — | 0.2724 | — |
| rs148110342 | R81C | 0.024 | 0.008 | 0.034 | 1.6E-03 | 0 | 1.6E-04 | 0.0015 | 0.4468 | 0.0005 | 0.0005 | 0.0055 | 0.0010 | 0.0005 | 0.0005 | 0.0005 | — | 0.0005 | — |
| rs751909359 | R86Q | 0 | 0.001 | 0 | 3.0E-05 | 0 | 0 | — | — | — | — | — | 0.0645 | — | — | — | — | — | — |
| rs752639409 | R637Q | 0 | 0.001 | 0 | 0 | 0 | 6.0E-05 | — | — | — | — | — | — | 0.1709 | — | — | — | — | — |
| rs761541008 | R677W | 0 | 0.001 | 0 | 5.0E-05 | 0 | 0 | — | — | — | — | — | 0.1000 | — | — | — | — | — | — |
| rs35641241 | R745Q | 0 | 0 | 0 | 1.2E-03 | 1.0E-03 | 8.0E-05 | — | — | — | — | — | — | — | — | — | 0.5857 | 0.0005 | 0.0005 |
| rs193268286 | S685P | 0 | 0 | 0 | 4.0E-05 | 0 | 0 | — | — | — | — | — | — | — | — | — | — | — | — |

The missense variants were defined as deleterious when predicted to be damaging, probably damaging, disease causing and deleterious by the five algorithms applied (SIFT, PolyPhen-2 HumVar, PolyPhen-2 HumDiv, MutationTaster and LRT score) and/or CADD score of more than 20. We considered only deleterious variants with minor allele frequency less than 1% in the burden analysis. P values are calculated using Chi-square test. KWT-Kuwaitis; IRN-Iranians; QAR-Qataris; EUR-Europeans (non-Finnish); EAS-East Asians; AFR-Africans.

## Discussion

At the time of writing this study, SARS-CoV-2 has infected 4,271,689 people worldwide, with over 1,665,097 cases in Europe, and 1,385,834 cases in the United States, approximately 82,919 cases in China, 109,286 cases in Iran, 23,623 cases in Qatar and 9286 cases in Kuwait ^1^. Mortality rates varied significantly around the world, with the highest reported deaths per 1 million people reported in Belgium, Spain, Italy, United Kingdom, and France reaching 756, 572, 508, 472, and 408 deaths/1 million respectively ^1^. Deaths per 1 million people in China, Japan, Qatar, Kuwait, and South Africa were significantly lower at 3, 5, 5, 15, and 3 respectively and moderately higher in Iran that reported the highest deaths per 1 million people in Asia at 80 deaths/1 million people ^1^. The lower infection and fatality rates in developing countries obviously cannot be attributed to increased number of tests or better medical services than what is already provided in Europe. Moreover, prior knowledge of the SARS-CoV-2 RNA genome revealed by whole genome sequencing of the virus, performed by us and others, has shown remarkable genetic similarities among the virus ^40^ circulating between Qatar, Kuwait and Europe, which was attributed to traveling and repatriation between these countries (Manuscript in preparation; GISAID ^40^). For these reasons, we argued that the most likely explanation for the differences observed in mortality rate among countries may be attributed to genetic variation in human genes involved in SARS-CoV-2 processing and cellular entry or exit. Therefore, we screened the genetic variations and eQTL expression of the SARS-CoV-2 candidate genes, *ACE2*, *TMPRSS2* and *FURIN* in three Middle Eastern populations: Kuwaiti, Iranian, and Qatari and compared them to available MAF data in the gnomAD database ^41^.

In this study we showed that amongst the nine known *ACE2* missense variants, N720D (rs41303171) and K26R (rs4646116) were the most frequent in the global datasets ^18^. In agreement with our analysis, structural predictions by Stawiski and colleagues revealed that the K26R missense variant enhanced the affinity of *ACE2* for SARS-CoV-2 whereas N720D had little involvement in the SARS-CoV-2 S-protein interaction ^18^. Our data suggest that the *ACE2* receptor N720D variant may enhance *TMPRSS2* activation and subsequent viral entry. Interestingly, the prevalence of the most *ACE2* activating variant N720D (rs41303171) was highest among Europeans (2.5%), Iranians (0.6%) when compared to Kuwaitis (0.3%), Qataris (0.2%) and other global populations (0.4%) and MAF of these variants significantly correlated with mortality rates in the corresponding countries and lower infection rates in Kuwait (Fig. 3). However, further functional assessments are required to confirm our predictions.

Additionally, we showed high prevalence of frequent deleterious *FURIN* variants in the Middle Eastern populations, especially Kuwait and Qatar, which indicated decreased *FURIN* protease activity that further reduces the risk of viral infection. As such, our results suggest a possible protective role of *FURIN* gene variants against the SARS-CoV-2 infection in the studied Middle Eastern populations. However, the data presented here add another variable that needs to be urgently addressed. It is not unreasonable to suggest that European human genetic variation presents optimal access for SARS-CoV-2 infection, whereas Iranian genetic background presents sub-optimal opportunities for the infection, and finally, Kuwaitis and Qataris have the least natural inducive genetic background in the *ACE2* and *FURIN* proteins combined for viral entry.

While the MAF of *ACE2* variant, R708W in Kuwait is 0.105%, which may indicate again a protective role in the Kuwaiti population, the absence of this and the other four deleterious variants from Africa, East-Asia ^41^, Qatar ^42^ does not explain the lower disease burden in these countries. Similarly, we did not observe a significant difference in the burden of novel deleterious variants comparing the Kuwaiti population with Europeans (Table 2), although they may play a protective role against COVID-19 locally. The functional roles of the two Kuwait-specific and the two Iran-specific *ACE2* deleterious novel variants Q661P (MAF=0.1%), A372G (MAF=0.2%) and V485L (MAF=0.18%), F452V (MAF=0.08%) respectively are yet to be experimentally determined.

Notably, the *ACE2* novel variant Q661P is close to the aa region 652-659, which is important for cleavage by the metalloprotease ADAM17. R716H and the two deleterious variants R708W and R708Q are located within residues 697-716 essential for cleavage by *TMPRSS11D* and *TMPRSS2*. In fact, the mutation of arginine-such as R708- and lysine residues within aa residues 697-716 markedly reduced *ACE2* cleavage by *TMPRSS2* ^43^. However, it should be mentioned that sequentially distant aa residues in the *ACE2* receptor can be seen brought structurally proximal to each other to create active sites for catalysis ^28^. We illustrated this in Supplementary Fig. 6 to show that the novel aa changes (colored green) are proximal to the protein residues that mediate its activity (colored blue and red). Further studies are needed to directly assess the functional aspects of the reported missense variants. We urge the international community to assess *ACE2* variation differences among people with mild/asymptomatic disease versus patients presenting with severe respiratory distress syndrome.

Interestingly, there was not a single individual in the Middle East with *ACE2* receptor variations in amino acids known to be crucial for SARS-CoV-2 S-protein binding (K31, E35, D38, M82, K353) ^44^. This may indicate that natural immunity conferred by the *ACE2* receptor variations is lacking or extremely rare. The rareness is evident from a recent comprehensive study that scrutinized *ACE2* gene variations in more than 290,000 individuals from 400 different worldwide population groups and in which only eight rare *ACE2* gene missense variants (K31R, E35K, E37K, D38V, N33I, H34R, Y83H, and Q388L) with reduced binding to the S-protein of SARS-CoV-2 were reported ^18^. Notably, this study has disproved the claim of Cao and colleagues on the absence of such protective *ACE2* variants in human populations ^15^.

Finally, according to the Human Protein Atlas portal, the mRNA and protein expression of the *ACE2* gene is predominant in the human testis, cardiovascular and type II pneumocytes (https://www.proteinatlas.org/ENSG00000130234-ACE2/tissue). Further, a recent study that profiled scRNA-seq from the human testicles revealed the presence of this receptor in Spermatogonia, Leydig and Sertoli Cells ^45^. These findings may indicate that human testis is a potential target for coronavirus conforming to male prevalence in infected cases. Notably, in our data, Kuwaiti individuals carrying *ACE2* missense variants were all males. Also, a Chinese population study observed a higher hemizygous mutation rate in males than females ^15^. It is also possible that the high hemizygosity seen in males can be responsible for male prevalence in infected cases. This hypothesis can be further supported by the recent detection of SARS-CoV-2 virus in the semen of infected people ^46,47^.

In summary, we report the differential occurrence of gene variants in the Middle Eastern populations that enrich the knowledge on the genetic susceptibility of different global populations to the novel SARS-CoV-2 virus. Especially that in the context of a pandemic, rare variants that increase or decrease disease susceptibility become quantitatively important since millions of people may be infected. The decreased burden of rare deleterious variants and the increased MAF in variants that activate the *ACE2* receptor in the European population suggest an inherent susceptibility to SARS-CoV-2 infection and *vice versa* in the Middle Eastern population. We understand that other confounding factors like SARS-CoV-2 testing, socioeconomical status, the availability of proper medical services and the higher burden of other diseases may be important contributors to the disparities seen in mortality rates around the world. It is imperative that more comprehensive studies should be conducted as more patient data emerges from different parts of the world. Our data are preliminary and highlight the urgent need to correlate patients’ medical/infection history to their genetic variants in order to predict in a more accurate and personalized way how human genetic variations may influence this pandemic.

## Methods

### Genetic data

The whole-exome sequences data of 473 Kuwaitis ^48^, 800 Iranians ^49^ and 100 Qataris ^42^ published previously were used in this analysis. The genetic data of non-Finnish European, East Asian and African American populations were obtained from the gnomAD repository ^41^, which contain data on a total of 125,748 exomes and 71,702 genomes (https://gnomad.broadinstitute.org/).

### Gene expression data

The expression data for *ACE2*, *TMPRSS2,* and *FURIN* were obtained from the genotype-tissue expression (GTEx) database (https://gtexportal.org/home/). The same database portal was used to extract quantitative trait loci (eQTLs) for the three genes.

### Statistical analysis

The missense variants were defined as deleterious when predicted to be damaging, probably damaging, disease causing and deleterious by the five algorithms applied, SIFT ^50^, PolyPhen-2 HumVar, PolyPhen-2 HumDiv ^51^, MutationTaster ^52^ and LRT score ^53^ and/or CADD (Combined Annotation-Dependent Depletion) score of more than 20 ^54^. We considered only deleterious variants with minor allele frequency (MAF) less than 1% in the burden analysis. The significance of the differences in MAFs between different populations was calculated using Chi-Square test, using the R software (https://www.r-project.org/). All the p-values presented in the tables are not corrected for multiple testing. P values ≤0.05 were considered significant.

### Structural Analysis

All the identified *ACE2* missense exon variants were mapped, modeled, and analyzed using Pymol modeling software (https://pymol.org/2/). DynaMut web server was used to predict the effect of genetic variants on the stability and flexibility of *ACE2* receptor ^32,33^.

## Acknowledgements

This work was supported by Coronavirus emergency resilience grant from Kuwait Foundation for the Advancement of Sciences.

## Author contributions

F.A-M. conceptualized the study. A.M., and A.A.M. conducted structural predictions and modeling. D.H. created figures. M.E. organized the data and conducted most analyses. T.A.T., S.E.J., and A.C., collected the Middle Eastern populations’ data. R.N. performed whole-exome sequencing. F.A-M., and M.E. wrote the manuscript. H.A., M.A-F., R.A., and J.A. critically appraised and edited the manuscript. All authors discussed, critically read and revised the manuscript, and gave final approval for publication.

## Materials & Correspondence

Correspondence and requests for materials should be addressed to F.A-M.

## Supplementary Information

**Supplementary Figure 1:** The positions of the *ACE2* receptor polymorphisms on the linearized *ACE2* protein. The translated protein contains an N-terminal signal sequence (1-17), single catalytic domain (18-740) with zinc-binding motif (HEMGH 374-378), a transmembrane region (741-761), a small C-terminal cytosolic domain (762-805).

**Supplementary Figure 2:** The positions of the *ACE2* receptor polymorphisms on 3D-model of the *ACE2* protein. 3D-structure superposition of polymorphic variants mapped to human *ACE2* monomer (Cyan) with full length structure of human *ACE2* (PDB ID: 6M18, Green). Active side zinc-binding residues (Magenta). Zinc atom (Pink). Key amino acids (Blue) and their prospective variants (Yellow).

**Supplementary Figure 3:** Structural superposition of polymorphic variants mapped to human *ACE2* with two native structural conformations of human *ACE2*. For both conformational structures, the SARS-CoV-2 Binding Domain is depicted in Red color, the active site Zinc Binding Domain is colored Blue. **(a)** Superposition of *ACE2* open conformational structure, PDB ID: 1r41 (Green) and the mapped model with the variants (Yellow). The identified SNPs (Magenta) at *ACE2* surface and within the molecule (Insert) depict conformational changes to original wildtype represented by the observed superimposed Yellow color. **(b)** Superposition of *ACE2* closed conformational structure PDB ID: 1r42 (Green) with a small peptide substrate shown in orange and the modeled structure (Yellow). The identified SNPs (Magenta) at *ACE2* surface and within the molecule (Insert) depict conformational changes to original wildtype represented by the observed superimposed yellow color.

**Supplementary Figure 4:** The positions of the *TMPRSS2* polymorphisms on the linearized *TMPRSS2* protein. The bar plot shows the frequencies of the selected variants that fall within critical *TMPRSS2* domains.

**Supplementary Figure 5:** The positions of the *FURIN* polymorphisms on the linearized *FURIN* protein.

**Supplementary Figure 6**. *ACE2* protein structure (open form, yellow; and closed form with substrate, gray). The active site residues are coded in red color. The zinc binding residues are coded in blue color. The identified novel amino acid changes in the Middle Eastern populations are coded in green color. The novel changes are proximal to the protein residues that mediate its activity.

**Supplementary Table 1**. *ACE2* eQTL variants in the Middle East and gnomAD populations.

**Supplementary Table 2**. *TMPRSS2* known missense variants in the Middle East and gnomAD populations.

**Supplementary Table 3.** Burden of *TMPRSS2* rare variants in the Middle East and gnomAD populations.

**Supplementary Table 4.** *TMPRSS2* eQTL variants in the Middle East and gnomAD populations.

**Supplementary Table 5.** *FURIN* eQTL variants in the Middle East and gnomAD populations.

