## Supplementary Information for "A comprehensive germline variant and expression analyses of *ACE2*, *TMPRSS2* and SARS-CoV-2 activator *FURIN* genes from the Middle East: Combating SARS-CoV-2 with precision medicine"

**Supplementary Table 3.** Burden of *TMPRSS2* rare variants in the Middle East and gnomAD populations.

**Supplementary Table 4.** *TMPRSS2* eQTL variants in the Middle East and gnomAD populations.

**Supplementary Table 5.** *FURIN* eQTL variants in the Middle East and gnomAD populations.

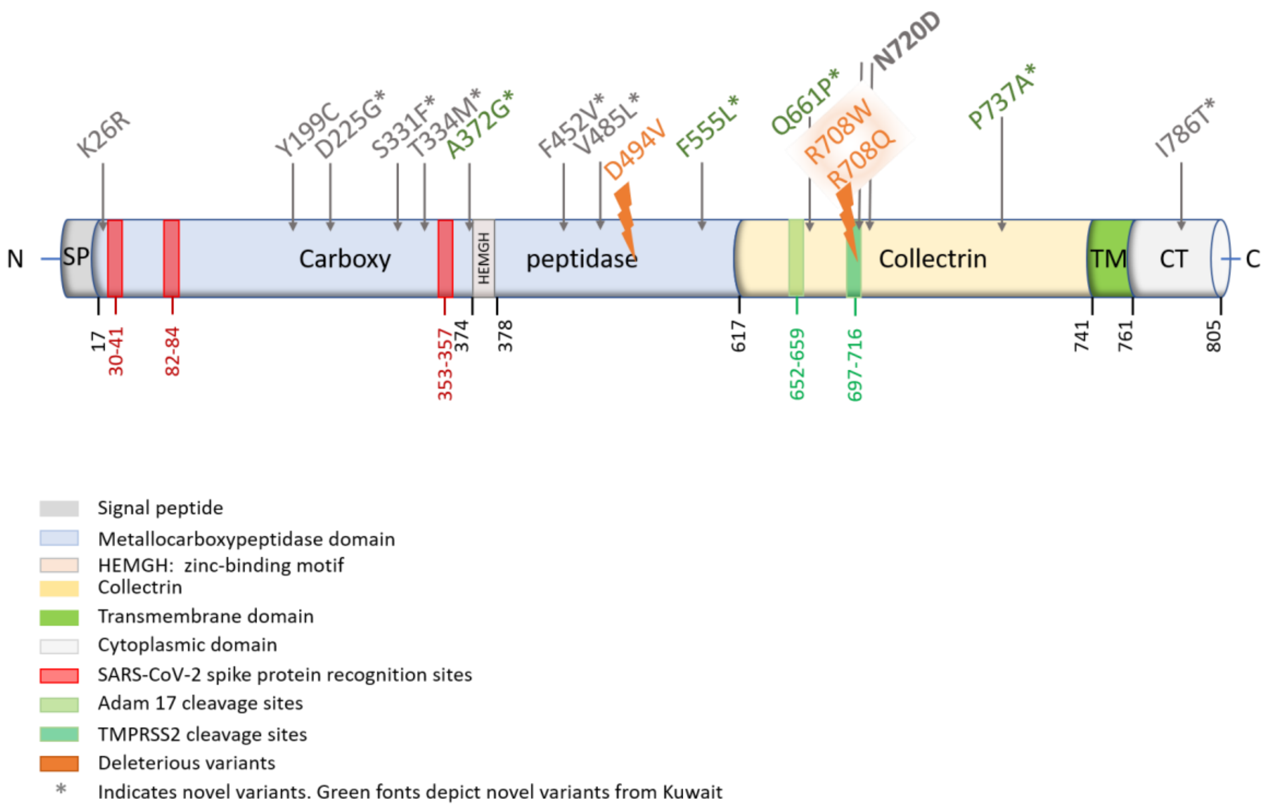


Supplementary Figure 1. The positions of the *ACE2* receptor polymorphisms on the linearized *ACE2* protein. The translated protein contains an N-terminal signal sequence (1-17), single catalytic domain (18-740) with zinc-binding motif (HEMGH 374-378), a transmembrane region (741-761), a small C-terminal cytosolic domain (762-805).

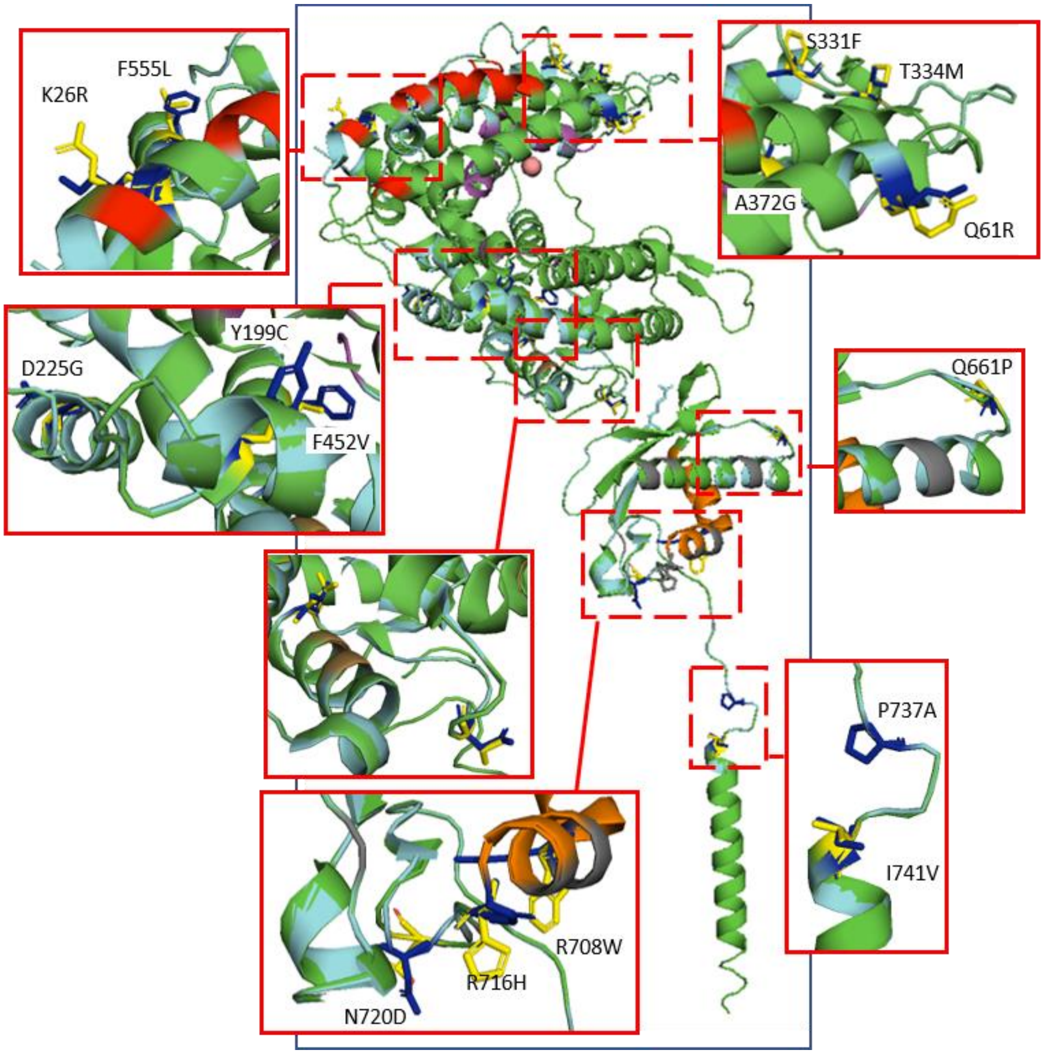


Supplementary Figure 2. The positions of the *ACE2* receptor polymorphisms on 3D-model of the *ACE2* protein. 3D-structure superposition of polymorphic variants mapped to human *ACE2* monomer (Cyan) with full length structure of human *ACE2* (PDB ID: 6M18, Green). Active side zinc-binding residues (Magenta). Zinc atom (Pink). Key amino acids (Blue) and their prospective variants (Yellow).

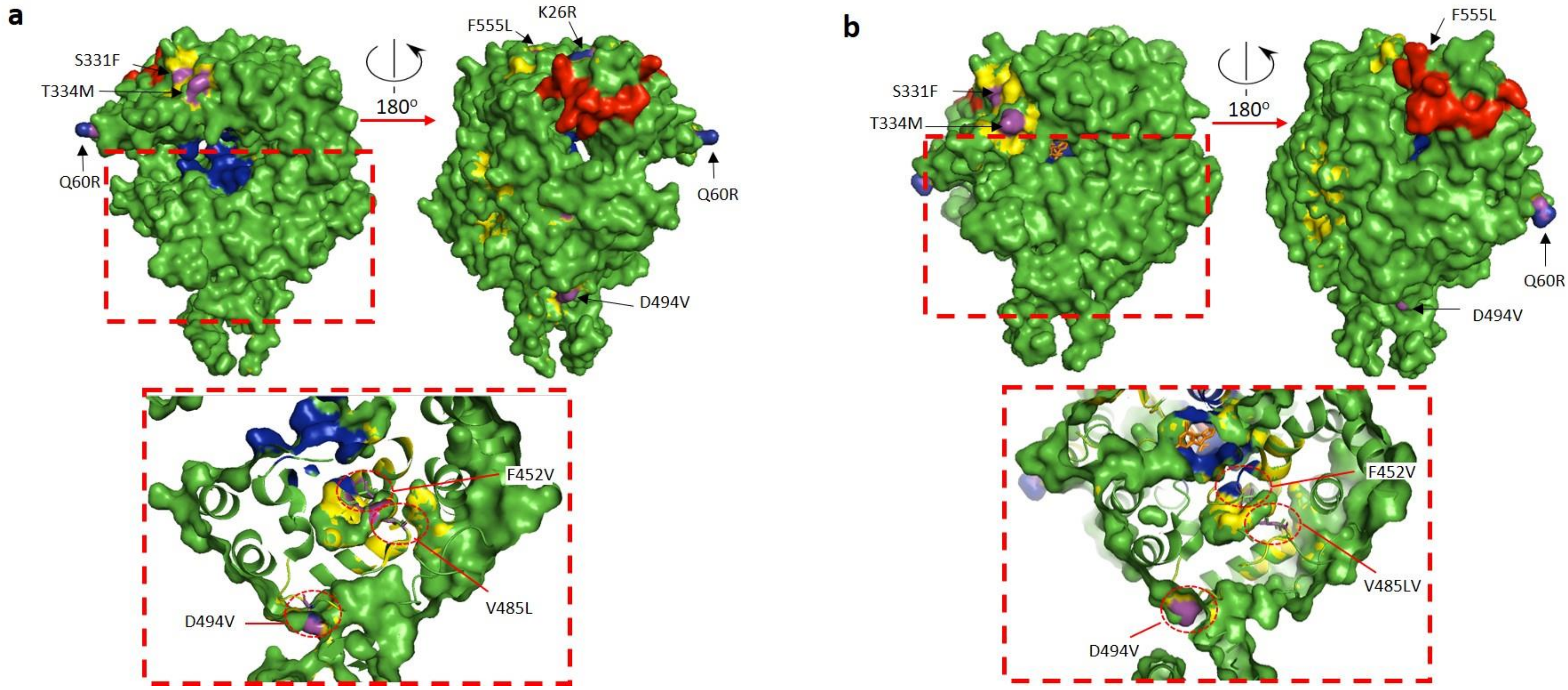


**Supplementary Figure 3.** Structural superposition of polymorphic variants mapped to human *ACE2* with two native structural conformations of human *ACE2*. For both conformational structures, the SARS-CoV-2 Binding Domain is depicted in Red color, the active site Zinc Binding Domain is colored Blue. **(a)** Superposition of *ACE2* open conformational structure, PDB ID: 1r41 (Green) and the mapped model with the variants (Yellow). The identified SNPs (Magenta) at *ACE2* surface and within the molecule (Insert) depict conformational changes to original wildtype represented by the observed superimposed Yellow color. **(b)** Superposition of *ACE2* closed conformational structure PDB ID: 1r42 (Green) with a small peptide substrate shown in orange and the modeled structure (Yellow). The identified SNPs (Magenta) at *ACE2* surface and within the molecule (Insert) depict conformational changes to original wildtype represented by the observed superimposed yellow color.

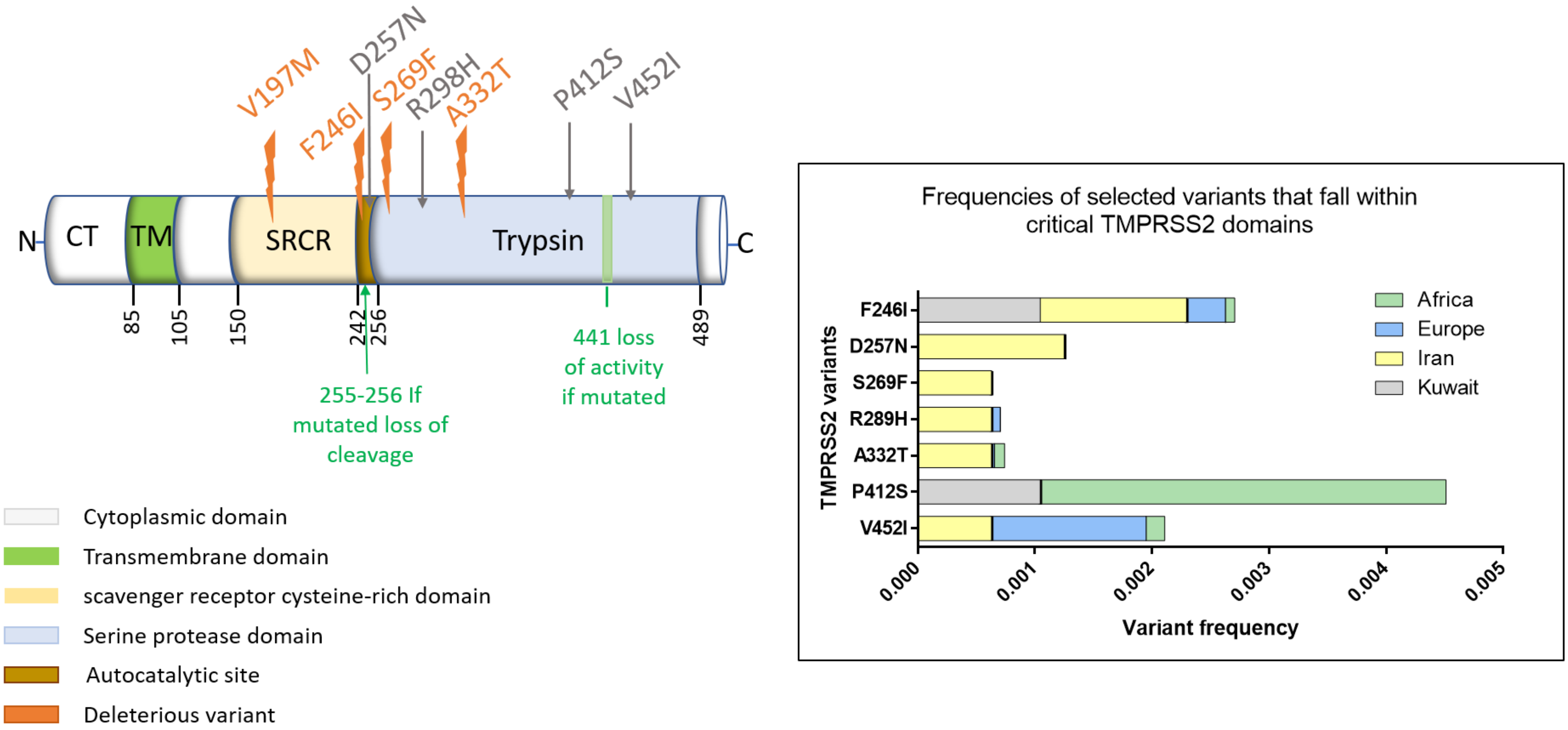


**Supplementary Figure 4.** The positions of the *TMPRSS2* polymorphisms on the linearized *TMPRSS2* protein. The bar plot shows the frequencies of the selected variants that fall within critical *TMPRSS2* domains.

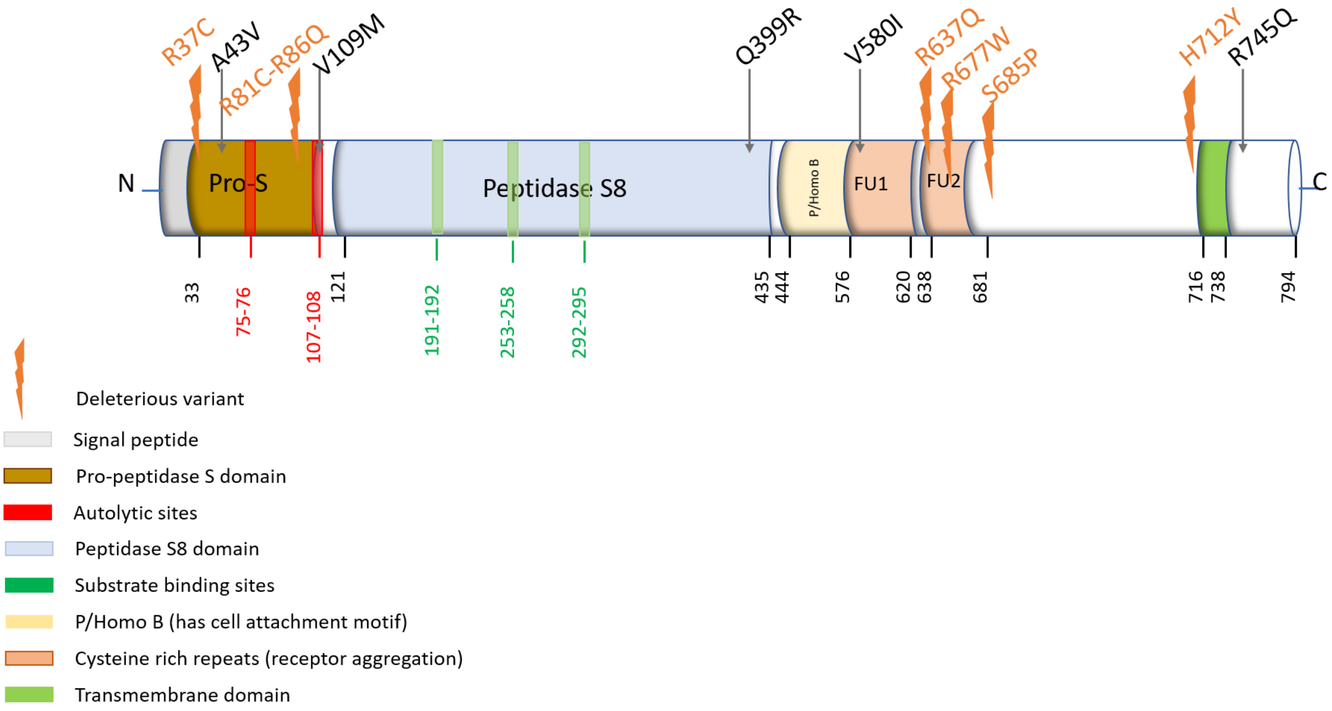


**Supplementary Figure 5.** The positions of the *FURIN* polymorphisms on the linearized *FURIN* protein.

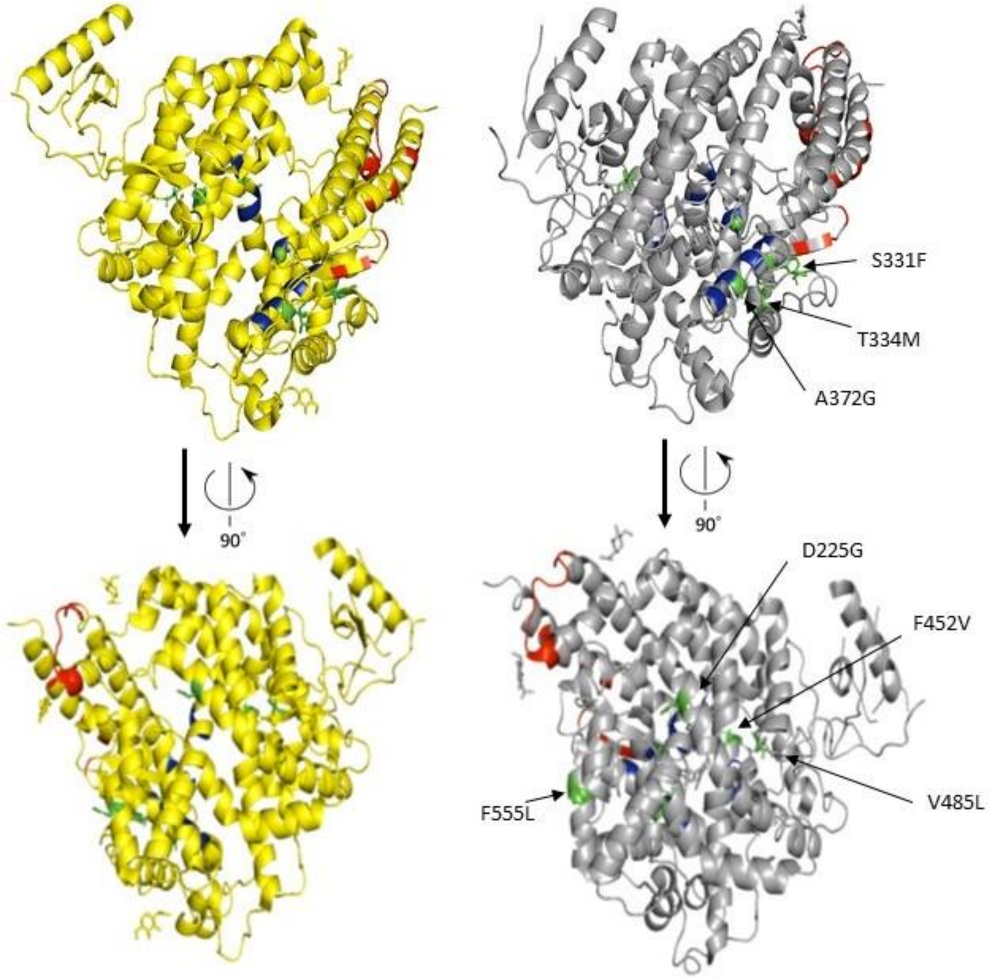


**Supplementary Figure 6.** *ACE2* protein structure (open form, yellow; and closed form with substrate, gray). The active site residues are coded in red color. The zinc binding residues are coded in blue color. The identified novel amino acid changes in the Middle Eastern populations are coded in green color. The novel changes are proximal to the protein residues that mediate its activity.

**Supplementary Table 1.** ***ACE2*** **eQTL variants in the Middle East and gnomAD populations.**

| Variant ID | Gene | Tissue | P GTEx | NES GTEx | Minor Allele Frequency | | | | | |
|---|---|---|---|---|---|---|---|---|---|---|
| | | | | | KWT | IRN | QAR | EUR | EAS | AFR |
| rs112171234 | *ACE2, BMX* | Adipose – Visceral (Omentum) | 4.1E-05 | -0.85 | NA | NA | 0.06 | 6.4E-04 | 0.00 | 0.20 |
| rs75979613 | *FANCB* | Breast - Mammary Tissue | 3.3E-05 | -0.44 | NA | NA | 0.01 | 0.01 | 0.00 | 1.7E-03 |
| rs12010448 | *MOSPD2, ASB9* | Muscle - Skeletal | 9.8E-05 | 0.43 | NA | NA | 0.03 | 1.9E-04 | 0.00 | 0.11 |
| rs4646127 | *ACE2* | Nerve - Tibial | 7.5E-09 | 0.2 | NA | NA | 0.58 | 0.62 | 1.00 | 0.78 |
| rs5936029 | *CA5BP1-CA5B* | Brain - Nucleus accumbens (basal ganglia) | 9.6E-16 | 0.6 | NA | NA | 0.49 | 0.48 | 1.00 | 0.87 |
| rs6632704 | *CLTRN* | Nerve - Tibial | 4.6E-16 | 0.26 | NA | NA | 0.41 | 0.48 | 0.95 | 0.50 |
| rs1996225 | *CA5BP1-CA5B* | Nerve - Tibial | 3.3E-17 | 0.27 | NA | 0.42 | 0.39 | 0.36 | 0.82 | 0.60 |
| rs6629110 | *ACE2* | Nerve - Tibial | 3.4E-15 | 0.25 | NA | NA | 0.41 | 0.47 | 0.96 | 0.43 |
| rs2158082 | *ACE2* | Nerve - Tibial | 1.0E-16 | 0.28 | 0.51 | NA | 0.55 | 0.48 | 1.00 | 0.93 |
| rs4830974 | *CLTRN* | Brain - Frontal Cortex (BA9) | 2.6E-17 | 0.60 | 0.47 | NA | 0.48 | 0.48 | 0.95 | 0.72 |
| rs5936011 | *CLTRN* | Brain - Nucleus accumbens (basal ganglia) | 3.2E-16 | 0.62 | NA | NA | 0.50 | 0.48 | 1.00 | 0.86 |
| rs4060 | *CA5BP1-CA5B* | Brain - Frontal Cortex (BA9) | 2.6E-17 | 0.60 | 0.04 | 0.50 | 0.44 | 0.48 | 0.95 | 0.65 |
| rs4830983 | *CA5BP1-CA5B* | Nerve - Tibial | 3.0E-16 | 0.28 | NA | NA | 0.51 | 0.48 | 1.00 | 0.93 |

P GTEx- P values reported in GTEx database; NES-normalized effect size in GTEx database (- sign/red font indicates down-regulation);
KWT-Kuwaitis; IRN-Iranians; QAR-Qataris; EUR-Europeans (non-Finnish); EAS-East Asians; AFR-Africans. NA denotes Not Available in exome data.

**Supplementary Table 2.** ***TMPRSS2*** **known missense variants in the Middle East and gnomAD populations.**

| **Variant ID** | **Protein Consequence** | **Minor Allele Frequency** | | | | | | **Functional Risk Prediction (Scores)** | | | | | |
|---|---|---|---|---|---|---|---|---|---|---|---|---|---|
| | | **Kuwait** | **Iran** | **Qatar** | **Europe** | **East Asia** | **Africa** | **SIFT** | **PP2HVAR** | **PP2HDIV** | **MUTTASTER** | **LRT** | **CADD** |
| rs148125094 | V452I | 0 | 0.00063 | 0 | 0.00132 | 0 | 0.00016 | T(0.28) | B(0.098) | B(0.018) | N(0.021) | N(0.111) | 7.914 |
| rs61735795 | P412S | 0.00105 | 0 | 0 | 0.00001 | 0 | 0.00346 | T(0.31) | B(0.086) | B(0.097) | N(0.043) | N(0.0025) | 12.16 |
| rs372563970 | A332T | 0 | 0.00063 | 0 | 0.00003 | 0 | 0.00009 | D(0.00) | D(0.97) | D(0.999) | D(0.898) | D(0.000) | 25.2 |
| rs775404304 | R289H | 0 | 0.00063 | 0 | 0.00007 | 0 | 0 | T(0.20) | B(0.01) | B(0.023) | N(0.459) | N(0.476) | 10.10 |
| rs547544037 | S269F | 0 | 0.00063 | 0 | 0 | 0 | 0 | T(0.72) | B(0.009) | B(0.007) | D(0.769) | N(0.125) | 22.1 |
| rs777951933 | D257N | 0 | 0.00125 | 0 | 0.00002 | 0 | 0 | T(0.55) | B(0.041) | B(0.101) | D(0.986) | N(0.005) | 15.17 |
| rs150554820 | F246I | 0.00105 | 0.00125 | 0 | 0.00033 | 0 | 0.00008 | T(0.27) | P(0.549) | P(0.935) | D(0.933) | N(0.022) | 22.5 |
| rs12329760 | V197M | 0.18200 | 0.19560 | 0.24279 | 0.23200 | 0.3838 | 0.29180 | D(0.02) | D(0.938) | D(0.999) | P(0.807) | N(0.022) | 24.8 |
| rs199865751 | D158N | 0 | 0 | 0.00119 | 0.00003 | 0 | 0 | T(0.47) | B(0.051) | B(0.251) | N(0.002) | N(0.009) | 14.77 |
| rs190265904 | L132F | 0 | 0.00188 | 0 | 0.00002 | 0.0004 | 0 | T(0.50) | D(0.943) | D(0.996) | N(0.035) | N(0.007) | 19.96 |
| rs147711290 | L128Q | 0 | 0 | 0.00568 | 0 | 0 | 0.00633 | T(0.07) | D(0.957) | D(0.999) | N(0.136) | N(0.302) | 23 |
| rs61735793 | T112I | 0.00628 | 0.01125 | 0.00469 | 0.01060 | 0 | 0.00169 | T(0.20) | B(0.015) | B(0.061) | N(0.005) | N(0.292) | 8.751 |
| rs770214639 | T95M | 0 | 0.00063 | 0 | 0.00002 | 0.00015 | 0 | T(0.05) | D(0.987) | D(1.00) | N(0.106) | D(0.0001) | 22.3 |
| rs141232947 | V73A | 0 | 0 | 0.00117 | 0 | 0 | 0.00024 | T(0.84) | B(0.002) | B(0.00) | N(0.0002) | N(0.709) | 0.002 |
| rs749665029 | A65V | 0.00105 | 0 | 0 | 0.00003 | 0 | 0 | T(0.54) | B(0.164) | B(0.452) | N(0.009) | N(0.531) | 2.407 |
| rs61735791 | A65T | 0.00209 | 0.00125 | 0 | 0.00283 | 0.00115 | 0.00088 | T(0.49) | B(0.029) | B(0.231) | N(0.007) | N(0.531) | 0.252 |
| rs61735790 | H55R | 0.00105 | 0 | 0.00781 | 0.00002 | 0.00005 | 0.00946 | T(0.30) | B(0.033) | B(0.074) | D(0.597) | N(0.008) | 15.96 |
| rs75603675 | G8D | 0.08159 | 0.33355 | 0.36364 | 0.42470 | 0.01298 | 0.32840 | T(0.35) | B(0.319) | P(0.815) | N(0.004) | N(0.291) | 12.01 |

SIFT (Sorting Intolerant From Tolerant): D = damaging, T = tolerated;
PP2HVAR (PolyPhen-2 Polymorphism Phenotyping v2 HumVar): D = probably damaging, P = possibly damaging, B = benign;
PP2HDIV (PolyPhen-2 Polymorphism Phenotyping v2 HumDiv): D = probably damaging, P = possibly damaging, B = benign;
MUTTASTER (MutationTaster): A = disease causing automatic, D = disease causing, N = polymorphism, P = polymorphism automatic;
LRT (Likelihood Ratio Test): D = deleterious, N = Neutral, U = unknown;
CADD - Combined Annotation Dependent Depletion based scores.
The missense variants were defined as deleterious when predicted to be damaging, probably damaging, disease causing and deleterious by the five algorithms applied (SIFT, PolyPhen-2 HumVar, PolyPhen-2 HumDiv, MutationTaster and LRT score) and/or CADD score of more than 20. We considered only deleterious variants with minor allele frequency less than 1% in the burden analysis.

**Supplementary Table 3. Burden of *TMPRSS2* rare variants in the Middle East and gnomAD populations.**

| Variant ID | Protein Consequence | Minor Allele Frequency | | | | | |
|---|---|---|---|---|---|---|---|
| | | KWT | IRN | QAR | EUR | EAS | AFR |
| rs372563970 | A332T | 0.00 | 6.3E-04 | 0.00 | 3.0E-05 | 0.00 | 8.7E-05 |
| rs150554820 | F246I | 1.0E-03 | 1.3E-03 | 0.00 | 3.3E-04 | 0.00 | 8.0E-05 |
| rs147711290 | L128Q | 0.00 | 0.00 | 0.01 | 0.00 | 0.00 | 0.01 |
| rs770214639 | T95M | 0.00 | 6.3E-04 | 0.00 | 2.3E-05 | 1.5E-04 | 0.00 |
| rs547544037 | S269F | 0.00 | 6.3E-04 | 0.00 | 0.00 | 0.00 | 0.00 |

The missense variants were defined as deleterious when predicted to be damaging, probably damaging, disease causing and deleterious by the five algorithms applied (SIFT, PolyPhen-2 HumVar, PolyPhen-2 HumDiv, MutationTaster and LRT score) and/or CADD score of more than 20. We considered only deleterious variants with minor allele frequency less than 1% in the burden analysis.

KWT-Kuwaitis; IRN-Iranians; QAR-Qataris; EUR-Europeans (non-Finnish); EAS-East Asians; AFR-Africans.

**Supplementary Table 4.** ***TMPRSS2*** **eQTL variants in the Middle East and gnomAD populations.**

| Variant ID | Gene | Tissue | P GTEx | NES GTEx | Minor Allele Frequency | | | | | |
|---|---|---|---|---|---|---|---|---|---|---|
| | | | | | KWT | IRN | QAR | EUR | EAS | AFR |
| rs79391937 | *C2CD2* (intronic) | Thyroid | 8.2E-06 | -0.33 | NA | NA | 0.02 | 0.02 | 0.00 | 0.003 |
| rs6517673 | *TMPRSS2* (intronic) | Prostate | 1.8E-12 | -0.3 | NA | NA | 0.17 | 0.10 | 0.001 | 0.18 |
| rs79566442 | *DSCAM* (intronic) | Ovary | 1.4E-06 | -1.2 | NA | NA | 0.05 | 0.04 | 0.00 | 0.01 |
| rs11701542 | intergenic | Testis | 5.3E-23 | 0.6 | NA | NA | 0.30 | 0.35 | 0.32 | 0.35 |

P GTEx- P values reported in GTEx database; NES-normalized effect size in GTEx database (-sign denotes down-regulation).
KWT-Kuwaitis; IRN-Iranians; QAR-Qataris; EUR-Europeans (non-Finnish); EAS-East Asians; AFR-Africans. NA denotes Not Available in exome data.

**Supplementary Table 5.** ***FURIN*** **eQTL variants in the Middle East and gnomAD populations.**

| Variant ID | Gene | Tissue | P GTEx | NES GTEx | Minor Allele Frequency | | | | | |
|---|---|---|---|---|---|---|---|---|---|---|
| | | | | | KWT | IRN | QAR | EUR | EAS | AFR |
| rs117976310 | *S2VB, LOC101926928* | Adrenal Gland | 3.0E-06 | -1.1 | NA | NA | 0.01 | 0.02 | 0.00 | 0.002 |
| rs12904679 | *BLM, FURIN* | Esophagus - Mucosa | 3.1E-06 | -0.2 | NA | NA | 0.12 | 0.12 | 0.003 | 0.02 |
| rs4932373 | *FES* | Esophagus - Mucosa | 6.6E-19 | 0.28 | NA | NA | 0.30 | 0.33 | 0.09 | 0.13 |
| rs2238332 | *BLM* | Pituitary | 2.7E-06 | -0.17 | NA | NA | 0.36 | 0.36 | 0.33 | 0.70 |
| rs8039305* | *FURIN* | Esophagus - Mucosa | 1.7E-17 | 0.23 | NA | 0.41 | 0.49 | 0.48 | 0.17 | 0.81 |
| rs6226* | *FURIN* | Artery - Tibial | 1.6E-09 | 0.12 | 0.72 | 0.65 | 0.69 | 0.68 | 0.53 | 0.93 |
| rs12904055 | *CRTC3* | Brain - Putamen (basal ganglia) | 1.1E-05 | 0.2 | NA | NA | 0.24 | 0.22 | 0.17 | 0.31 |
| rs80160059 | *SV2B* | Colon - Sigmoid | 9.0E-06 | 0.54 | NA | NA | 0.06 | 0.001 | 0.00 | 0.11 |
| rs2071410 | *FURIN* | Esophagus - Mucosa | 1.7E-20 | 0.28 | NA | 0.26 | 0.27 | 0.33 | 0.06 | 0.18 |
| rs8026133 | *LOC101926928, SLCO3A1* | Minor Salivary Gland | 1.4E-06 | 0.83 | NA | NA | 0.12 | 0.01 | 0.09 | 0.47 |
| rs2227935 | *BLM* | Esophagus - Mucosa | 2.9E-10 | 0.32 | 0.05 | 0.03 | 0.03 | 0.07 | 0.0004 | 0.07 |
| rs16944923 | *BLM, FURIN* | Skin - Sun Exposed (Lower leg) | 9.0E-06 | 0.12 | NA | NA | 0.13 | 0.14 | 0.20 | 0.05 |
| rs28385078 | *BLM* | Esophagus - Mucosa | 5.7E-10 | 0.32 | NA | NA | 0.02 | 0.07 | 0.00 | 0.03 |
| rs116449376 | *BLM, FURIN* | Esophagus - Mucosa | 1.5E-09 | 0.34 | NA | NA | 0.02 | 0.06 | 0.00 | 0.04 |
| rs7495370 | *BLM, FURIN* | Whole Blood | 5.6E-07 | 0.072 | NA | NA | 0.64 | 0.57 | 0.78 | 0.47 |
| rs7165790 | *BLM* | Heart - Atrial Appendage | 4.4E-07 | -0.12 | NA | NA | 0.34 | 0.00 | 0.001 | 0.00 |

P GTEx- P values reported in GTEx database; NES-normalized effect size in GTEx database; *P values calculated using Chi-square test. KWT-Kuwaitis; IRN-Iranians; QAR-Qataris; EUR-Europeans (non-Finnish); EAS-East Asians; AFR-Africans. NA denotes Not Available in exome data.
